# Graphical and Interactive Spatial Proteomics Image Analysis Workflow

**DOI:** 10.1101/2025.05.23.655879

**Authors:** Pritpal Singh, Jocelyn H. Wright, Kimberly S. Smythe, Bryce Fukuda, Ling-Hong Hung, Cecilia CS Yeung, Ka Yee Yeung

## Abstract

Spatial proteomics provides a spatially resolved view of protein expression and localization within cells and tissues by mapping the location and abundance of proteins. There is a need for fully-integrated end-to-end imaging workflows for spatial proteomic analysis that are flexible, high-throughput, and support graphical and interactive visualizations. We present a modular and interactive spatial proteomic image analysis workflow with individual steps containerized that empowers biomedical researchers to reproducibly execute and customize complex analyses.

Our workflow consists of cell segmentation, unsupervised clustering, validation of clusters on the image, and cell type clustering results visualization. Users can utilize a form-based graphical interface to execute and customize multi-step workflows with a single click or interactively adjust image processing steps within the workflow, apply workflows to various datasets, and modify input parameters as needed. We illustrated the functionality of our workflow using a cancer imaging dataset consisting of a tissue microarray (TMA) stained by high-plex immunohistochemistry. This TMA contained a variety of cancer and tissue cell types to assess the broad applicability of this workflow to different biopsy and tissue types.

## Introduction

The advent of high-plex spatial proteomics allows interrogation of patient tissue samples with over fifty-five antibody protein markers simultaneously, allowing for novel spatial biomarkers (e.g., specific cell type interactions, protein marker co-expression, or specific morphologies) to be identified that can predict disease. Techniques such as **co-d**etection by ind**ex**ing (CODEX), run on the PhenoCycler-Fusion (PCF) platform, are currently among the leading spatial proteomics imaging approaches. Analyzing spatial proteomics imaging data requires multiple analysis steps and many of these steps require a graphical user interface to facilitate the interpretation of results. Analysis of protein staining must consider not only expression level of a biomarker as is the case with spatial transcriptomics, but also the location of the staining (nuclear, cytoplasmic, or membranous), the staining pattern (dot like, linear, contiguous, diffuse and homogenous), and integration of cell morphology to allow for correct interpretation. Moreover, as this approach relies on antibody staining, a quality control step is essential for data validation.

Numerous software platforms can be used to analyze spatial proteomics imaging data, but they often come with limitations that pose challenges for users. Multi-modal analytical workflows often consist of analysis steps performed using different analysis tools or components that are not interoperable. A key limitation of existing approaches is the lack of an *end-to-end* software solution that enables users to interact with graphical visualizations and provide domain knowledge to guide the analyses. Most workflow execution engines focus on automating, orchestrating, and optimizing the execution of complex tasks but often lack native support for graphical outputs. However, support for interactive graphical analysis is essential to facilitate interpretation of imaging data. Another desirable feature is the flexibility to integrate open-source tools that are widely adopted by the bioimage community. As an example, a commercial platform offered by Enable Medicine [1] offers graphical output, consistency and reproducibility of analysis when images are uploaded to their proprietary server. In addition to the lack of support for *on-prem* analysis, it is not possible to integrate widely used bioimage tools like QuPath [2] and data output is not flexible. As another example, SPEX (Spatial Expression Explorer) is an open-source modular analysis platform with a graphical user interface that supports spatial proteomics and spatial transcriptomics workflows [3]. However, the support for interactive analysis of graphical output and flexibility of integrating commonly used graphical tools are limited.

Here, we present a modular and interactive graphical platform that empowers biomedical researchers to reproducibly execute complex spatial proteomic analysis in an automated and end-to-end reproducible workflow that easily integrates multiple open-source software tools and custom scripts through containerization. Containerization enables reproducibility and portability by bundling a software application, along with all its dependencies and specific configurations, into a single file known as a container image. This allows the software to run consistently across various computing environments. Our spatial proteomics imaging workflow supports graphical output and is packaged in version-controlled software containers, ensuring reproducibility and interoperability. The end-to-end workflow consists of cell segmentation, unsupervised clustering, validation, and interactive graphical visualization of results. The workflow is customizable to allow changes in user defined analysis parameters or inclusion of additional analysis modalities for spatial analysis of cell types.

## Analytical Workflow and Case Study

We developed a spatial proteomics imaging workflow and applied it to a tissue microarray (TMA) stained by high-plex immunohistochemistry. The TMA contained both normal and tumor tissues consisting of common and antigenically diverse cancers including breast adenocarcinoma (BRC), non-small cell lung carcinoma (NSCLC), colorectal cancer (CRC), melanoma (MEL), hepatocellular carcinoma (HCC), and Merkel cell carcinoma (MCC). The images were stained using PCF. PCF offers deep insights into tissue architecture, cellular organization, and the role of the microenvironment in diseases like cancer. Previous analysis of PSF data using the open-source programs, QuPath [2] to visualize and segment the data and CytoMAP [4] for clustering analysis, were successful but cumbersome to move the data back and forth between the two programs and not always reproducible. Analysis on the Enable Medicine platform was faster and consistent, but data output was not flexible. In addition, analysis on Enable Medicine required images to be uploaded to their server which is time consuming and expensive to store image data and perform analyses. Most importantly, sharing data with third party servers complicates data management, compliance, privacy, and security issues.

Building on our previous experience working with CODEX data, we designed and developed a multi-step analytical workflow leveraging open-source programs and protocols (**Figure 1A**). This workflow is designed to allow customization by addition/modification of analysis steps, parameters, and output via modular containers. As outlined in **Figure 1A**, steps of the workflow include (1) visualization of the stained image for assessing quality control of individual stains, (2) segmentation of cells into cell data units, (3) exporting these data to perform unsupervised clustering of cell types, (4) creation of masks to be superimposed onto the image for visual validation, and (5) generation of data summaries such as heatmaps and dimension reduction techniques (e.g., UMAP) to facilitate cell-type annotation and distribution between samples.

**Figure 1:**
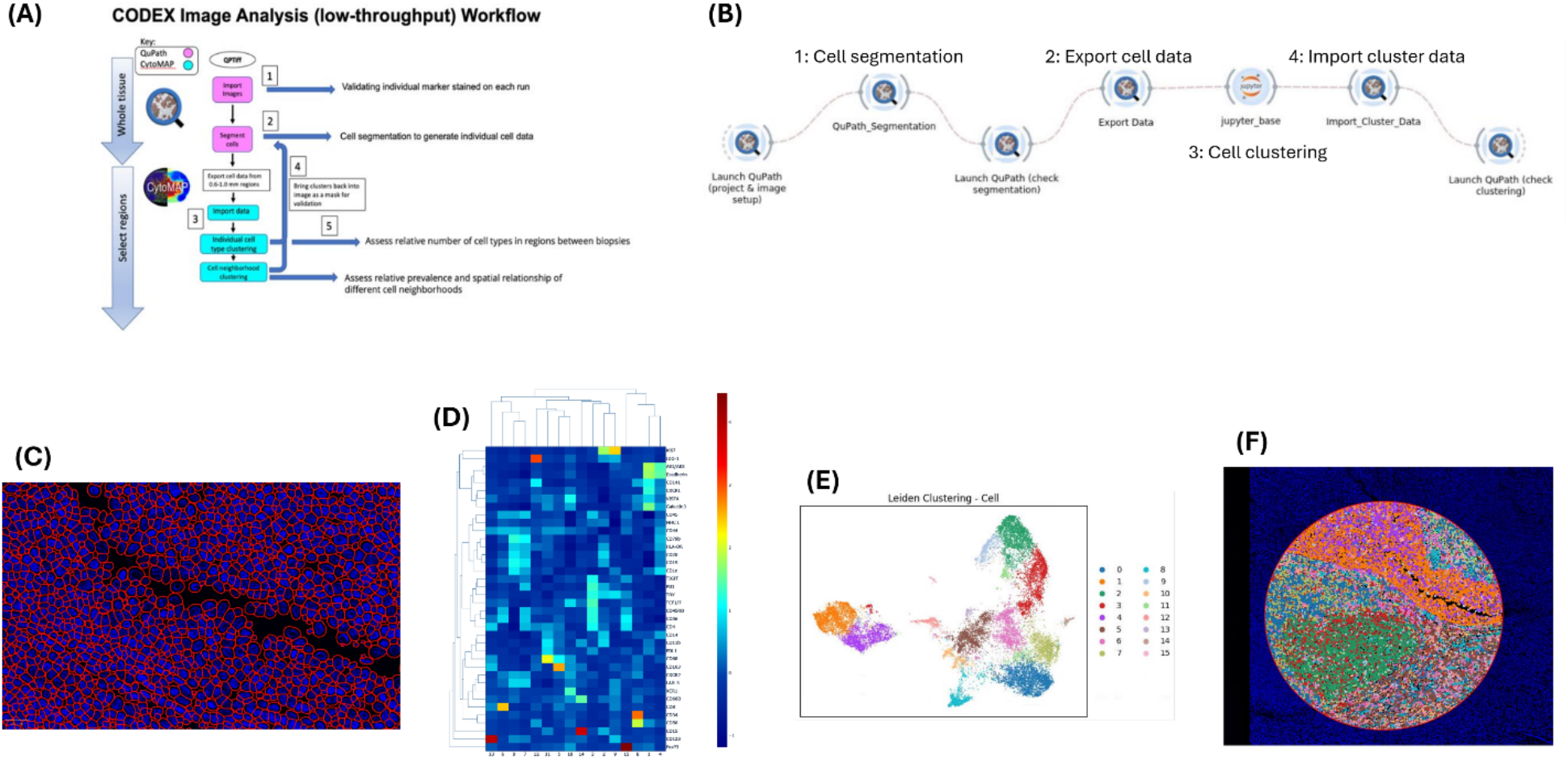
(A) A schematic diagram of a manual low-throughput analytical protocol for cell segmentation and unsupervised clustering using QuPath and CytoMAP image analysis platforms used at Fred Hutch is shown. (B) A screenshot showing the high-throughput graphical containerized workflow consisting of modular widgets. (C) This is an image in QuPath illustrating the output from our containerized workflow on a tonsil region from a stained TMA image. Results of cell segmentation are shown as red cell outlines surrounding DAPI (blue) stained cell nuclei that define the cell data objects. (D) and (E) show results of Leiden unsupervised clustering of exported cell data from the segmented image from C as a heatmap in D, and as a UMAP in E. (F) The results of importing clustering results from E back into QuPath and displaying it as a colormap over the original tonsil tissue image is shown.

## Implementation

We implemented a high-throughput automated workflow of analytical protocols adopted in our CLIA accredited laboratory at the Fred Hutchinson Cancer Center. Our spatial proteomics analysis workflow, as illustrated in **Figure 1B**, leverages the Biodepot-workflow-builder (Bwb) platform [5, 6], a desktop application that supports interactive graphical output and each task (or module) in the workflow is represented by a graphical widget. These widgets are containerized using Docker containers and linked to other widgets to produce a full workflow. Users can interact with each widget and adjust the parameters using a form-based user interface, and they can incorporate new tools and/or scripts into their workflow. See the Supporting Information for additional implementation details.

## Results

Our workflow allows running QuPath within a containerized environment to view and interact with image(s) and QC markers as if run as a standalone desktop application. Using the workflow, one can select regions on the image(s) and perform cell segmentation within a QuPath project using an optimized version of the StarDist algorithm [7] and export the cell object data containing x, y coordinates and mean cell fluorescence pixel intensity (MFI) values for each biomarker into a csv file. The exported cell data can then be used to perform unsupervised clustering of cell types using the Leiden algorithm and visualizing the clustering results as Uniform Manifold Approximation and Projections (UMAP) and heatmaps using Jupyter Notebooks and python libraries. Lastly, the workflow allows importing clustering data back into QuPath to make colormaps (masks) to review with the original image as an overlay to validate the results. All analysis steps in the workflow can be executed either end-to-end with a single click or run individually in an interactive manner.

Analysis was tested on our multi-cancer TMA, and it gave accurate cell segmentation and biologically relevant cell type clustering. **Figures 1C, 1D, 1E, and 1F** show example results of running cell segmentation, unsupervised clustering, and cluster colormap generation on a region of human tonsil through our containerized workflow. Following cluster validation on the image, parameters could be interactively modified using the graphical user interface shown in **Figure 1B**, and the analysis rerun, enabling reproducible and iterative analysis.

## Conclusions

Image analysis workflows often consist of multiple steps, each of which runs in a different software tool that in turn require different dependencies and computing environments. This approach leads to major challenges in reproducibility, interoperability, and scalability, due to lack of systematic version control and local hardware. Moreover, most image analyses contain steps that are graphical and require interactive user input, features that are not supported by most workflow execution engines. We addressed these challenges by developing an open-source, containerized, graphical, and cloud-enabled spatial proteomics analysis workflow that easily integrates with other software through containerization.

We integrated the QuPath image analysis tool as a containerized widget to automate our manual image analytical protocol consisting of cell segmentation, unsupervised clustering, and results visualization. Users can modify individual steps or widgets and corresponding input parameters using a drag-and-drop user interface. Additional modalities can easily be incorporated in this graphical workflow as additional widgets for specific types of spatial analysis desired, such as proximity values between individual cell types and cellular neighborhood analysis. Importantly, users can reproduce the work of others or incorporate additional analyses into their own workflows with ease. Finally, our workflows can be deployed locally, on internal computer servers or on the cloud such that data security and compliance can be directly managed by the user. This contrasts with commercial solutions in which sensitive patient data must be uploaded and analyzed on external proprietary server for which the user has limited control.

## Availability of Data and Software

Raw data associated with this paper are publicly available at Zenodo with DOI 10.5281/zenodo. 15825670 (URL: https://zenodo.org/records/15825670).

Source code and documentation of software are publicly available at https://github.com/BioDepot/SpatialProteomics

## ONLINE SUPPORTING INFORMATION SUPPLEMENTARY METHOD

### Creation of widgets and workflows in the Biodepot-workflow-builder

Our Bwb workflow automates a low-throughput manual workflow. There are various widgets you can interact with within this workflow to do your image analysis. You can interact with each widget individually or run them as part of your end-to-end workflow ad hoc. Each widget is dependent on the QuPath Dockerfile used to containerize it.

The “Launch_QuPath” widget is used to launch a standalone instance of QuPath within BwB for managing your QuPath project, performing your analysis, and visualizing the results. This widget has two optional parameters to specify that the QuPath project file and image open when QuPath is launched. If these parameters are not specified, QuPath will launch without a project open by default.

### Cell Segmentation

The segmentation widget allows you to run StarDist segmentation within QuPath on your entire image or an individual TMA core by specifying it as a parameter of the widget. StarDist is a deep learning-based image segmentation method specifically designed for biological images, such as those obtained from microscopy. This widget depends on my_stardist.groovy script and stardist_cell_seg_model.pb StarDist model file. The two required fields of this widget include qpProjFile and image_to_segment. Users can optionally specify image resolution and TMA core to segment via the image_resolution and core_to_segment parameters. By default, the widget will segment all the tissues in the specified image with image resolution set to 0.5. Default settings are tailored for PCF QPTIFF images, however parameters can be adjusted for analysis of Keyence QPTIFFS, as well as images with OME-TIFF format from Lunaphore COMET platforms.

### Clustering

The “jupyter_base” widget launches the BWBQuPathClustering.ipynb Jupyter Notebook that runs the Leiden unsupervised clustering algorithm Leiden on the QuPath cell data in the all-cell-measurements.csv file, exported via the “Export_Image_Data” widget. The cell data is normalized before clustering. The notebook then depicts the clustered data using UMAPs (Uniform Manifold Approximation and Projection) and heatmaps. Clusters in the UMAP plot can be used to identify groups of cells that belong to the same cell type based on their patterns of biomarker expression. The UMAP is generated using the Python scanpy package, utilizing the scanpy.pp.neighbors function to generate neighbors, with n_neighbors. The scanpy.tl.umap function is used to generate the UMAP plot with min_dist parameter. The default values for n_neighbors and min_dist are 30 and 0.0001 correspondingly but can be configured in the notebook. The interactive unsupervised clustering heatmap is plotted using the python graphing library Plotly and it uses rows to represent biomarkers and columns to represent clusters. The final output of the notebook is the clustering data, written to leiden_clustering_export.csv in the clustering_data_export directory.

### Overlay of clustering results with images

The “Import_Cluster_Data” widget imports the clustering data from the csv files in the clustering_data_export directory into QuPath to generate colormap overlays (masks) onto the original image using the import_clusters.groovy script. Each cluster is assigned a unique color. The spatial arrangement of these clusters is displayed directly on the tissue image which allows to see whether certain clusters localize to specific regions, providing insights into tissue architecture or pathology. The one and only required parameter for this widget is the qpProjFile to specify the QuPath project for importing the clustering data.

## ACKNOWLEDGEMENT

The flowchart in Figure 1A is inspired by materials from Akoya Biosciences [8]. We would also like to thank Grady Carson for personal communications and Mike Nelson for his discussion forum post on image.sc at https://forum.image.sc/t/there-and-back-again-qupath-cytomap-cluster-analysis/43352.

## DECLARATIONS

### Author’s Contributions

J.H.W., K.S.S., C.Y., L.H.H. and K.Y.Y. conceived and supervised the study. P.S. developed and tested the workflow and performed the analysis. B.F. tested and validated the workflow. J.H.W., K.S.S. and C.Y. provided the data and use cases. P.S., J.H.W., C.Y. and K.Y.Y. wrote the draft manuscript. K.S.S., B.F., L.H.H., and K.Y.Y. were involved in discussions and contributed to the manuscript. All authors read and approved the final manuscript.

